# Engineered extracellular vesicles targeting BACE1 reduces amyloid-beta plaque formation in a genetic mouse model of Alzheimer’s Disease

**DOI:** 10.64898/2026.08.10.744066

**Authors:** Valerie S. Kalluri, Sara Che, Meagan Conner, Barbara Moreno Diaz, Aman Yarlagadda, Kaira A. Church, Antonios Chronopoulos, Karina Vázquez-Arreguín, Hikaru Sugimoto, Raghu Kalluri

## Abstract

Alzheimer’s disease (AD) is a progressive neurodegenerative disorder characterized by the accumulation of amyloid-β (Aβ) plaques, neurodegeneration, and cognitive decline. β-Site amyloid precursor protein cleaving enzyme 1 (BACE1) catalyzes the rate-limiting step in Aβ production and remains a therapeutic target for AD. However, effective delivery of RNA therapeutics to the brain remains challenging due to the blood-brain barrier (BBB). Here, we evaluated the feasibility of using clinical-grade mesenchymal stem cell-derived extracellular vesicles (EVs) as systemic carriers for *Bace1*-targeting small interfering RNA (siRNA) in the 5xFAD mouse model of AD. Engineered EVs crossed the BBB and delivered siRNA cargo to the brain, with uptake observed in both neurons and astrocytes. Systemic therapy with EVs engineered to encapsulate Bace1 siRNA resulted in reduced brain Bace1 protein levels and a decrease in amyloid plaque burden compared with control EVs carrying scrambled siRNA. The reduction was most pronounced in larger, high-intensity plaques, suggesting that Bace1 suppression may preferentially limit plaque growth and maturation. Repeated systemic administration was well tolerated, with no evidence of treatment-associated toxicity. These findings establish a proof-of-concept feasibility for EV-mediated delivery of Bace1-targeting siRNA to the brain and support further development of engineered EVs as a therapeutic platform for neurodegenerative diseases. Future studies incorporating behavioral, molecular, and mechanistic analyses will be required to determine the extent to which Bace1 suppression delivered through EVs can modify disease progression and improve functional outcomes in AD.

## Introduction

Alzheimer’s disease (AD) is the most common cause of dementia and affects more than 55 million people worldwide. The disease is characterized by progressive cognitive decline, memory impairment, synaptic dysfunction, and neurodegeneration, ultimately leading to severe disability and death^1^. Neuropathologically, AD is defined by the accumulation of extracellular amyloid-β (Aβ) plaques and intracellular neurofibrillary tangles composed of hyperphosphorylated tau protein, accompanied by widespread neuroinflammation, neuronal loss, and disruption of neuronal networks^2^. Although AD is a multifactorial disorder, genetic, pathological, and biomarker evidence supports a central role for Aβ accumulation in disease initiation and progression, particularly during the preclinical stages of disease^3,4^.

Aβ peptides are generated through sequential proteolytic processing of amyloid precursor protein (APP). In the amyloidogenic pathway, APP is first cleaved by β-site APP-cleaving enzyme 1 (BACE1), producing a membrane-associated C-terminal fragment that is subsequently processed by γ-secretase to generate Aβ peptides, including the aggregation-prone Aβ40 and Aβ42 species^5^. Because BACE1 catalyzes the rate-limiting step in Aβ production, it has been regarded as an attractive therapeutic target for AD^6^. The importance of BACE1 in AD pathogenesis has been demonstrated through multiple genetic studies in transgenic mouse models. Complete deletion of Bace1 in APP-overexpressing mice abolished excess Aβ40 and Aβ42 production and restored cognitive performance to levels observed in wild-type animals^7^. Similarly, Bace1 deficiency in 5XFAD mice prevented Aβ accumulation and blocked the development of AD-like neuropathology^8^. Partial reduction of Bace1 expression also delayed disease progression in 5XFAD mice, resulting in significant reductions in amyloid plaque burden during early disease stages; however, these benefits diminished with age as BACE1 expression increased and pathology advanced^9,10^. Importantly, inducible deletion of BACE1 in adult 5XFAD mice reduced established amyloid pathology and improved cognitive function, demonstrating that therapeutic suppression of BACE1 can provide benefit even after disease onset^11^. Collectively, these studies provide evidence that BACE1 is a potential target for AD intervention.

Despite promising preclinical results in genetically engineered mice, translation of Bace1 inhibition to the clinic has proven challenging. Multiple BACE1 inhibitors, including verubecestat, atabecestat, and lanabecestat, effectively reduced Aβ production in humans but failed to demonstrate clinical benefit and were associated with adverse effects that led to discontinuation of development programs^12–14^. Emerging evidence suggests that these outcomes may reflect the diverse physiological functions of BACE1 beyond APP processing, including roles in synaptic plasticity, axonal guidance, and myelination^6,11^. While complete and sustained pharmacological inhibition of BACE1 may disrupt essential neuronal processes, this does not invalidate BACE1 as a therapeutic target and instead highlight the need for alternative approaches capable of achieving more controlled and selective suppression of BACE1 expression. By selectively reducing expression of disease-associated genes, small interfering RNAs (siRNAs) may permit tunable target suppression while minimizing off-target effects. Numerous studies have demonstrated the therapeutic potential of Bace1-targeted RNAi in AD models. Lentiviral delivery of Bace1-targeting RNAi to APP transgenic mice reduced Bace1 expression, attenuated amyloidogenic APP processing, decreased neurodegeneration, and improved spatial learning and memory^15^. Similarly, systemically administered PEGylated peptides-based nanocomplexes carrying Bace1 siRNA reduced Bace1 expression, decreased amyloid plaque burden, and improved cognitive performance in APP/PS1 mice^16^. Intravenously administered extracellular vesicles (EVs) engineered with a neuron-specific rabies virus glycoprotein (RVG) targeting peptide successfully crossed the blood-brain barrier (BBB), delivered Bace1 siRNA to the brain, and reduced cortical BACE1 expression in vivo^17^. Together, these studies establish proof-of-principle that RNA-based suppression of BACE1 can mitigate AD-associated pathology.

The clinical translation of RNA therapeutics for neurological disorders is limited by challenges associated with delivery, stability, biodistribution, and efficient transport across the BBB^17–20^. EVs/Exosomes have emerged as promising delivery vehicles capable of overcoming many of these barriers. EVs are nanoscale membrane-bound vesicles released by all cells and act as important mediators of intercellular communication through the transfer of proteins, lipids, messenger RNAs, microRNAs, and other bioactive cargo molecules^21,22^. EVs are naturally evolved transport systems and their low immunogenicity and ability to access tissues that are difficult to reach using conventional delivery platforms, including the central nervous system^22^, favors their clinical translation potential. We previously reported on engineered EVs that successfully delivered siRNA and short hairpin RNA cargo targeting oncogenic Kras in pancreatic cancer models and Myc in glioblastoma, resulting in gene silencing and therapeutic benefit in mice^23–25^.

Mesenchymal stromal/stem cells (MSCs) represent an especially attractive platform for EV-mediated therapeutic delivery. Bone marrow derived MSCs produce abundant EVs that retain many of the biological properties of their parent cells and have been investigated as delivery vehicles for proteins, nucleic acids, and other therapeutic cargoes^22,25^. We recently reported on the safety profile of bone marrow derived EVs engineered to deliver therapeutic siRNA payload against oncogene Kras^G12D^ in patients with advanced pancreatic cancer^26^. Here we evaluated EVs bone marrow-derived MSCs loaded with Bace1-targeting siRNA through electroporation and administered to 5XFAD mice, a widely used transgenic model of AD^27^. In this proof-of-concept study, we aimed to determine the feasibility of MSC-mediated delivery of Bace1-targeting siRNA and to establish a foundation for the future development of EVs-based RNA therapeutics for Alzheimer’s disease.

## Methods

### Cell Culture

Neuro2A (CCL-131), 293T (CRL-3216), and SH-SY5Y (CRL-2266) cells were purchased from ATCC. Human bone marrow derived MSC were obtained as previously described^24,25^. Neuro2A cells were cultured in EMEM and MSCs were cultured in αMEM with ribonucleosides/deoxyribonucleosides and L-glutamine (Corning). EMEM was supplemented with 10% FBS and αMEM supplemented with 2 U/mL heparin (Sigma), 1% NEAA (Gibco), and 5% PLT Max (EMD Millipore). 293T cells were cultured in DMEM supplemented with 10% FBS. All media was further supplemented with 1% Penicillin-Streptomycin (Corning). Cells were cultured at 37°C and 5% CO2. Neuro2A, 293T, and SH-SY5Y cell identities were validated using STR fingerprinting and all tested negative for mycoplasma.

### Generation of Bone Marrow Derived Mesenchymal Stem Cell-Derived EVs with siRNA

EVs were generated from human bone marrow derived MSCs as previously described^24^. Briefly, EVs were purified by filtration through a 0.2 µm filter and ultracentrifugation of the conditioned culture media. Size and concentration of EVs were determined by particle tracking analysis (NanoSight LM10, Nanosight v3.1; camera level:13, detect threshold:5). EVs were characterized by flow cytometry for EV and MSC markers as previously described (see also **Supplementary Methods**). To generate EVs with siRNA payload, 4E9 EVs and 100pmol siRNA were mixed together in PlasmaLyte (400 μL total), then electroporated using Biorad Gene Pulser XM (exponential setting 400V, 125uF and infinite resistance). Two distinct sequences of siRNA were obtained from Dharmacon: siRNAs against *BACE1* (sense strands, siRNA-1: GCUUUGUGGAGAUGGUGGAUU, siRNA-2: UGGACUGCAAGGAGUACAAUU) and matched scrambled control sequences (sense strand, scrbl siRNA-1: GGUGGAUGCGUAGUGGUAUUU, scrbl siRNA-2: GAAAGGGUCUACGAACUGAUU). For targeting studies, Cy-5-tagged siRNA was used.

### In vitro BACE1 Targeting

Cells were treated with siRNA using transfection with Lipofectamine 2000 (20pmol siRNA) or engineered EVs (∼20-25pmol siRNA). Cells were collected 24 hours after treatment, and Direct-Zol (Zymo Research, Cat. No. R2050/2052) was used to extract RNA, followed by cDNA synthesis (Thermo Fisher, Cat. No. 4374966). *BACE1* expression was determined using qPCR with Quant Studio 7 (Thermo). Primers are listed in the **Supplementary Methods**. Technical replicates were averaged and relative expression expressed as fold change, with statistical analyses carried out on ΔCt when comparing biological replicates, defined as independent experiments.

### Animal Model and Treatment

5xFAD (B6SJL) female and wild-type (WT) mice were purchased from the Jackson Laboratory (JAX #6554). There is no information available on parental origin of transgene. The mice were allowed to acclimate in standard housing conditions at UT MD Anderson Cancer Center (MDACC) animal facilities for at least one week before experiment started. All procedures were reviewed and approved by Institutional Animal Care and Use Committee of MDACC. Ten weeks old female 5xFAD or WT mice were treated with EVs containing siRNA targeting Bace1 (iExo Bace1 siRNA) or scrambled siRNA (iExo Scrbl siRNA), with each treatment of 100 μL of 10^9^ EVs/exosomes containing approximately 25pmol siRNA (via electroporation, see above). For in vivo studies, siRNA-2 and matched scrambled sequence listed above were used. Treatment schedule was 3x per week (approximately every 2-3 days) via retro-orbital injection (intravenous, IV). The animals were euthanized after ∼13 weeks of treatment (41 treatments). Brains were collected for downstream analyses, including snap frozen in liquid nitrogen, processed for formalin fixation and paraffin embedding, or fixed in 4% PFA then 30% sucrose for histological analysis.

### Immunohistochemistry and immunofluorescent labeling studies

For immunohistochemistry (IHC) and immunofluorescent (IF) staining, a series of 5 µm FFPE sections were collected. To analyze for Aβ plaques, three sections (50 µm apart) were stained with 1% thioflavin T (Millipore Sigma, Cat. No. T3516) for 8 minutes, then destained with 80% and 95% ethanol. The frontal lobe was imaged (5-6 images/section, 10x objective) using Zeiss LSM800confocal microscope. To stain for Aβ, three sections (50 µm apart) were stained with 4G8 (Biolegend, Cat. No. 800701) and 3,3′- Diaminobenzidine was used as a chromogen. Images along the frontal lobe (5-6 images/section, 10x objective) as well as the hippocampus (representative images) were taken using Leica DM1000 LED microscope mounted with a Leica DFC295 microscope camera, Leica LAS version 4.4 software. For quantification, images were analyzed using FIJI sotfware. Number, size, and intensity of individual plaques (stained by ThT or anti-Aβ antibody) were analyzed using a macro and predetermined threshold values. The plaque area and integrated density for each mouse was used for comparison between groups. Quartile distributions of integrated density or Ab plaque area were also represented across all field of views. Frozen brain sections were also blocked with with 10% Donkey Serum with 0.5% Triton X and immunolabeled with for NeuN (Milllipore Sigma, Cat. No. MAB377, 1:4,000) or GFAP (Invitrogen, Cat. No. 130300, 1:4,000), and using nuclear counterstained with Hoechst (1:1,000). Z-stack Images (20x objective) were taken using Zeiss LSM800confocal microscope, and maximum projection image was used to evaluate colocalization of Cy5-tagged siRNA with NeuN and GFAP signal.

### Digital western blot

Proteins were extracted from frozen brain pieces using RIPA (ThermoScientific, Cat. No. 89900) with protease (Roche, Cat. No. 11697498001) and phosphatase (Roche, Cat. No. 4906845001) inhibitor by homogenization (Qiagen PowerLyzer 24 Homogenizer). Protein concentration was quantified using bicinchoninic acid (BCA) assay (Pierce, Cat. No. 23225). Digital western blot (200 ng protein) was performed using Wes (ProteinSimple) with *BACE1* (Cell Signaling Technology, Cat. No. 5606, 1:50) and β-actin (Cell Signaling Technology, Cat. No. 3700, 1:10,000) antibodies. Full length lanes are shown in **Supplementary Figure 1.**

### Statistical analysis

All statistics were analyzed using GraphPad Prism 10, and all tests used are defined in the figure legends. n values are listed and defined in the figure legends.

## Results

### Generation of EVs targeting BACE1

We defined two siRNA sequences targeting both murine and human Bace1 transcript, namely siRNA-1 and siRNA-2 (**Figure 1A**, see methods). The siRNA used for biodistribution was also labeled with Cy5 fluorescent tag (**Figure 1B**). EVs/exosomes were electroporated with siRNA to encapsulate siRNA inside EVs/exosomes (**Figure 1B**), as previously described^24^. Human bone marrow MSC EVs size was ascertained by NanoSight^TM^ tracking analysis. The size distribution showed a mean size of 139 nm and a mode size of 98 nm (**Figure 1C**). Flow cytometry analyses showed EVs display expression of MSC markers CD90 and CD29 and tetraspanins associated with EVs, namely CD63, CD9 and CD81 (**Figure 1D, Supplementary Figure 1A**). Concordant with our previous findings^24^, MSC EVs also display expression of CD47 (**Figure 1D**), the ‘don’t eat me’ signal minimizing phagocytic clearance.

**Figure 1.**
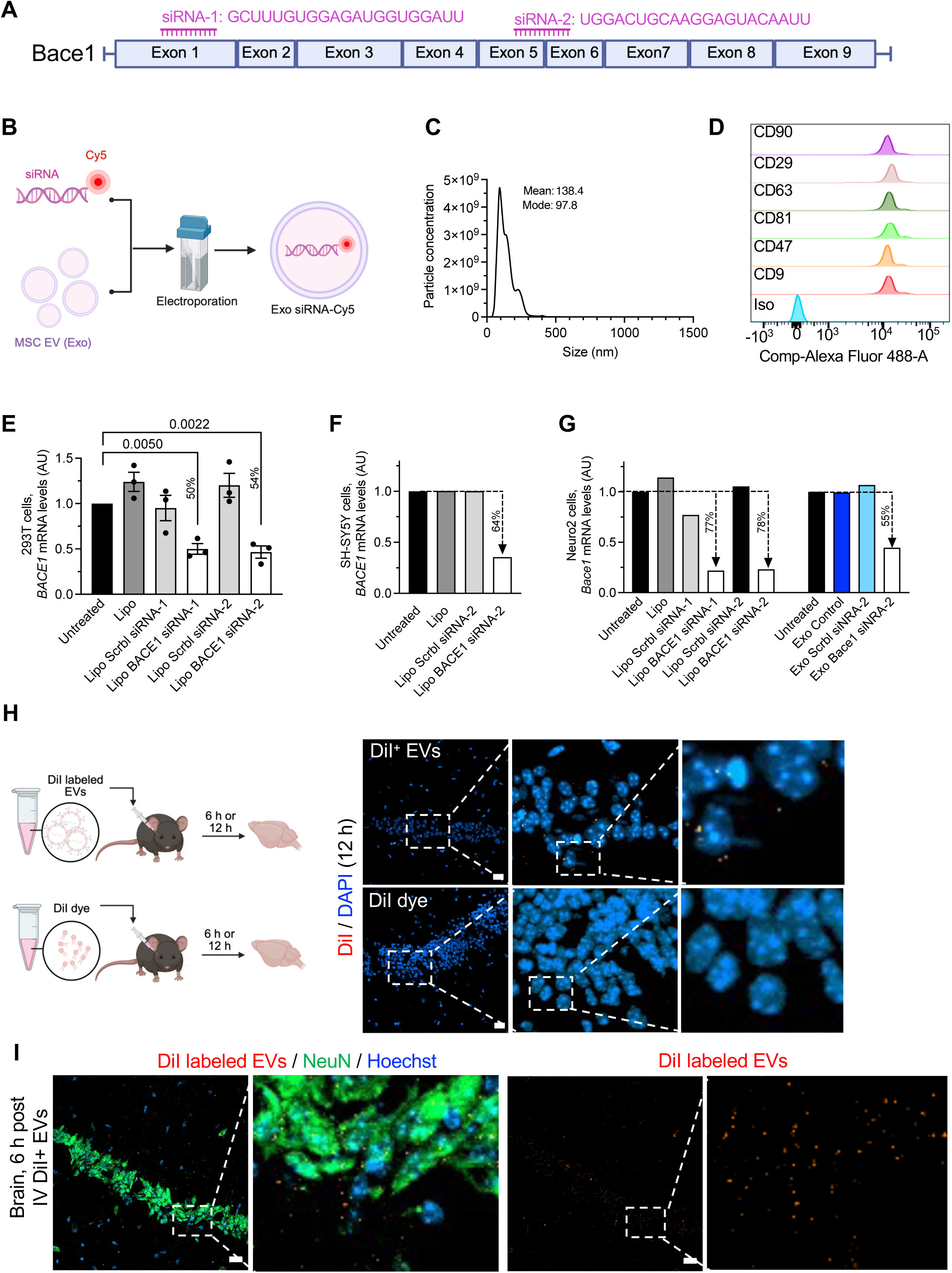
Generation of engineered EVs for brain delivery of Bace1 siRNA payload. **A**. Schematic representation of murine *Bace1* mRNA and siRNA sequences used to suppress Bace1 expression. Exon box sizes approximate the relative size of each exon. **B**. Schematic representation of the electroporation of bone marrow derived MSC EVs (Exosomes/Exo) to incorporate Cy5-tagged siRNA. **C**. Representative size distribution and particle concentration (per mL, using NanoSight^TM^). Mode and mean size in nm listed. **D**. Histogram depicting positive expression by flow cytometry of EVs for the listed markers in relation to isotype (Iso) control. See also **Supplementary Figure 1A** for gating strategy. **E**. Relative expression of *BACE1* in 293T cells following lipofectamine (Lipo) transfection of siRNA targeting *BACE1* or matched scrambled control sequence, n = 3 distinct experiments. One way ANOVA on ΔCt, p values are listed. **F**. Relative expression of *BACE1* in SH-SY5Y cells following lipofectamine transfection of siRNA targeting BACE1 or matched scrambled control sequence, n = 1 experiment. **G**. Relative expression of *Bace1* in Neuro2 cells following lipofectamine transfection or EV (Exo) treatment with siRNA targeting Bace1 or matched scrambled control sequence, n = 1 experiment. **H**. Schematic representation of the biodistribution experiment evaluating DiI labeled EVs compared to DiI dye alone control at 6 and 12 hours post intravenous administration of EVs/Dye. Representative image of brain section (hippocampal region) for DiI and nuclear stain (DAPI) with serial digital zooms at 12 hours post DiI^+^ EVs (top) or Dye only (bottom) administration. Scale bar: 20 μm. See also **Supplementary Figure 2** for individual fluorescent channels. **I.** Representative image of brain section (hippocampal region) for DiI, NeuN, and nuclear stain (Hoechst) with digital zoom at 6 hours post EV administration (Left). Single channel for DiI labeled EVs also shown (Right). Scale bar: 20 μm. See also **Supplementary Figure 3** for individual fluorescent channels. In **E-G**: percentages indicated relative decrease in transcript levels.

siRNA sequences targeting BACE1 in 293T cells were tested using lipofectamine transfection. Compared to untreated cells, lipofectamine alone (Lipo), or scrambled siRNA transfected (Scrbl) cells, 293T cells transfected with either siRNA sequence tested (BACE1 siRNA-1 or siRNA-2) show a specific decrease in BACE1 transcript level (**Figure 1E**). Specific decrease in BACE1 transcript levels is also seen in human immortalized neuroblastoma SH-SY5Y cells and murine neuroblastoma Neuro-2 cells transfected with BACE1 siRNA-2 (**Figure 1F-G**). Neuro-2 cells treated with EVs with Bace1 siRNA-2 (Exo Bace1 siRNA-2) show a 55% reduction in Bace1 transcript levels compared to control EV/exosomes (Exo Control, not electroporated) or EV with scrambled siRNA (Exo Scrbl siRNA-2) (**Figure 1G**). Taken together these results support the specific targeting of Bace1 in vitro in both murine and human cells with EV/Exo Bace1 siRNA, with siRNA-2 used for subsequent studies.

### Intravenously administered EVs localized to the brain

We previously reported that EVs administered intravenously, intraperitoneally, and intranasally localize to the brain in mice and non-human primates^23–26^. DiI labeled EVs are detected in the brain (hippocampus area) of mice 6 and 12 hours following intravenous (IV) injection, visualized by fluorescence microcopy of brain sections with DAPI nuclear stain (**Figure 1H, Supplementary Figure 2**). DiI^+^ EVs co-localizes with and without NeuN^+^ neurons in the hippocampus (**Figure 1I**, **Supplementary Figure 3A-B**). 5xFAD and littermate control (WT) mice were treated with Exo Bace1 siRNA or Scrbl siRNA, with 3x per week dosing IV over a period of 13 weeks (**Figure 2A**). Body weight measurements remain stable across all experimental groups, supporting that serial IV administration of engineered EVs are well tolerated (**Figure 2B-C**). Examination of brain tissue following IV administration of Exo Bace1 siRNA-2 with Cy5 fluorescent tag showed accumulation adjacent to DAPI stained nuclei (**Figure 2D**). Cy5 labeled Bace1 siRNA co-localized with NeuN expressing neurons (**Figure 2E**) and GFAP expressing astrocytes (**Figure 2F**).

**Figure 2.**
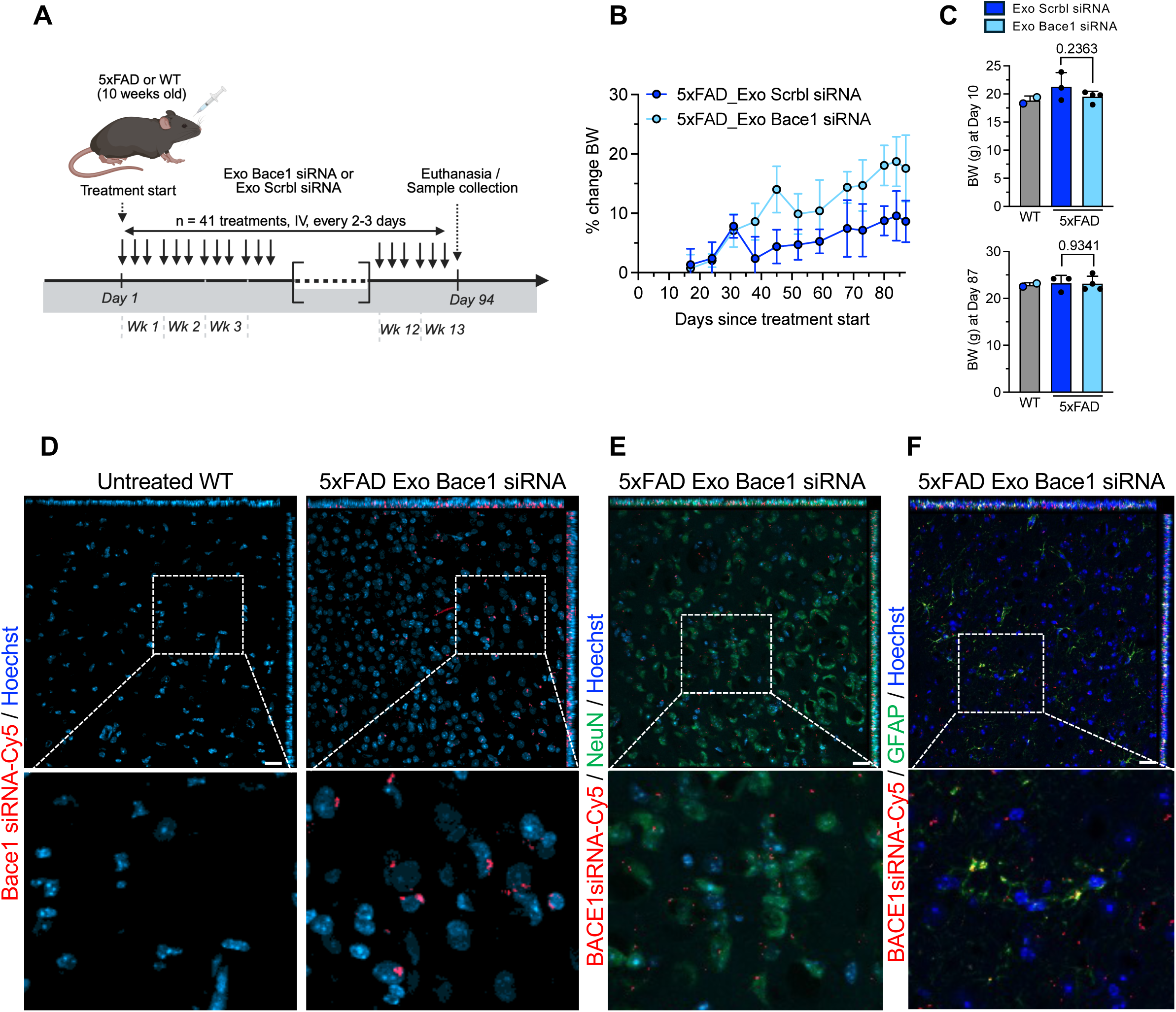
IV administered EVs with BACE1 siRNA cargo localizes to the brain. **A**. Schematic representation of experimental design. Control wild type (WT) or 5xFAD mice were treated with intravenous iExo Bace1 siRNA or control iExo Scrbl siRNA for 41 consecutive dosing approximately 2-3 days apart for a total of 41 doses over the course 13.4 weeks or 94 days. **B**. Body weight (BW) overtime expressed as percent change in the indicated groups. Exo Bace1 siRNA: n = 4 mice; Exo Scrbl siRNA: n =3 mice. **C.** Body weight at day 10 and day 87 post treatment in the indicated groups. WT control: n = 2 mice; Exo Bace1 siRNA: n = 4 mice; Exo Scrbl siRNA: n =3 mice. Unpaired two-tailed t test, p values are listed. **D.** Representative images of the brain of untreated wild type (WT) control mice and 5xFAD mice following Exo Bace1 siRNA with Cy5 tag (Bace1 siRNA-Cy5) with nuclear (Hoechst) counter stain. Scale bar: 20 μm. Digital zoom shown. **E.** Representative images of the brain of 5xFAD mice following Exo Bace1 siRNA with Cy5 tag (Bace1 siRNA-Cy5) with NeuN colocalization and nuclear (Hoechst) counter stain. Scale bar: 20 μm. Digital zoom shown. **F.** Representative images of the brain of 5xFAD mice following Exo Bace1 siRNA with Cy5 tag (Bace1 siRNA-Cy5) with GFAP colocalization and nuclear (Hoechst) counter stain. Scale bar: 20 μm. Digital zoom shown.

### EV with siRNA targeting BACE1 suppress plaque formation in 5xFAD mice

WT control and 5xFAD mice were treated with Exo-BACE1 siRNAor Exo-Scrbl siRNA as described above (**Figure 2A**) and euthanized at the study endpoint. Brains were harvested and stained with thioflavin T (ThT) to detect amyloid fibrils (**Figure 3A**). A reduced number of plaques per field of view was observed in Exo Bace1 siRNA-treated 5xFAD mice compared with Exo Scrbl siRNA-treated controls, based on analysis of fields of view from n = 3 Exo Scrbl siRNA-treated and n = 4 Exo Bace1 siRNA-treated mice (**Figure 3B**). Immunolabeling of the brains for beta amyloid (abnormally processed and precursor) showed a decrease in large plaques with high intensity (75% quartile), with minimal impact on smaller plaques (25% quartile) (**Figure 3C-D**). Across field of views, a decrease in plaque area and in integrated density is observed in mice treated with Exo Bace1 siRNA compared to Exo Scrbl siRNA (**Figure 3E**). A relative decrease in Bace1 protein levels is also observed in the brain of mice treated with Exo Bace1 siRNA compared to Exo Scrbl siRNA (**Figure 3F, Supplementary Figure 1B**).

**Figure 3.**
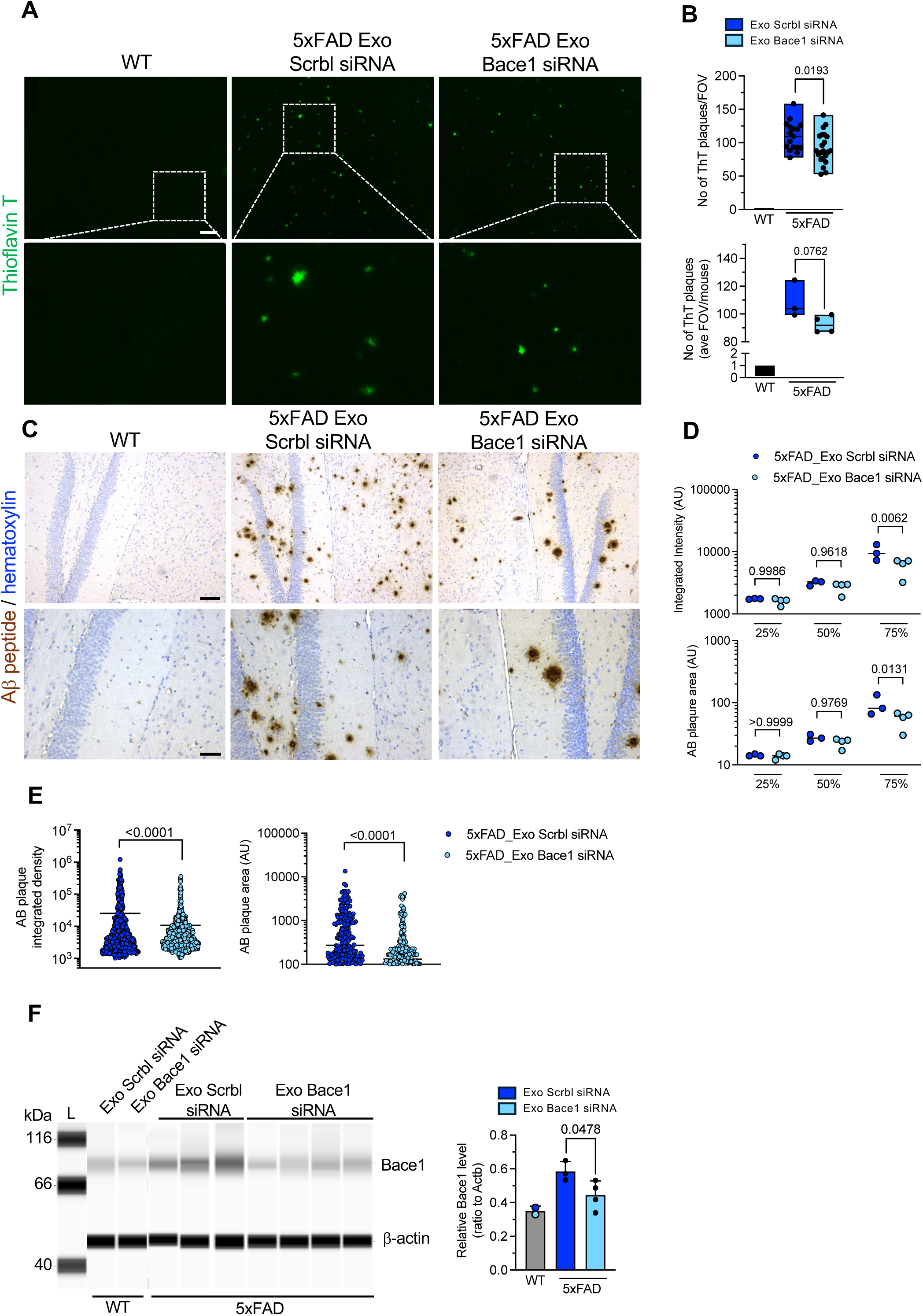
Exo BACE1 siRNA suppresses BACE1 and plaque formation in 5xFAD mouse brain. **A**. Representative images of Thioflavin T in the brain of WT control and 5xFAD mice following Exo Bace1 siRNA or Exo Scrbl siRNA control. Scale bar: 100 μm. Digital zoom shown. **B.** Quantification of the number of Thioflavin T (ThT) plaques per field of view (FOV) (top, each dot represents an FOV) and averaged per mice (bottom, each dot represents a mouse), WT control: n = 2 mice; Exo Bace1 siRNA: n = 4 mice; Exo Scrbl siRNA: n =3 mice. **C**. Representative images of Aβ peptide in the brain of WT control and 5xFAD mice following Exo Bace1 siRNA or Exo Scrbl siRNA control. Scale bar: 100 μm (top) and 50 μm (bottom). **D**. Quantification of plaque integrated density and area averaged per mice and presented along quartile distribution (25%, 50%, 75%), WT control: n = 2 mice; Exo Bace1 siRNA: n = 4 mice; Exo Scrbl siRNA: n =3 mice. One way ANOVA, p values are listed. **E**. Quantitation of plaque integrated density and area presented for all mice and all field of views examined (each dot is a plaque). Unpaired two-tailed t test with Welch’s correction, p values are listed. **F.** Relative Bace1 protein level to actin control in brain lysates from the listed mice assayed by capillary electrophoresis and associated quantification, WT control: n = 2 mice; Exo Bace1 siRNA: n = 4 mice; Exo Scrbl siRNA: n =3 mice. Unpaired two-tailed t test with Welch’s correction, p value is listed. L: ladder. kDa: kilodalton. See also **Supplementary Figure 1B** for full length run of electrophoretic migration of proteins.

## Discussion

In this proof-of-concept study, we demonstrate the feasibility of using engineered mesenchymal stem cell-derived extracellular vesicles (EVs) as a systemic delivery platform for Bace1-targeting siRNA in a genetic mouse model of Alzheimer’s disease. Repeated intravenous administration of EVs loaded with Bace1 siRNA resulted in detectable delivery of cargo to the brain, localization within neurons and astrocytes, reduction of brain Bace1 protein levels, and a corresponding decrease in amyloid plaque burden. Long-term administration over 14 weeks was well tolerated, with no overt evidence of toxicity as assessed by body weight stability throughout the treatment period. Collectively, these findings support the feasibility of EV-mediated RNA interference as a strategy to modulate pathogenic pathways within the central nervous system.

A major challenge for RNA therapeutics targeting neurodegenerative diseases is efficient delivery across the blood-brain barrier. While several nanoparticle and viral approaches have demonstrated proof-of-principle activity, limitations related to biodistribution, immunogenicity, and clinical scalability have limited clinical translation. Here we show that fluorescently labeled EVs and EV-encapsulated siRNA were detected within the brain parenchyma following systemic administration, with evidence of uptake by both NeuN-positive neurons and GFAP-positive astrocytes. These observations are consistent with previous reports demonstrating the intrinsic ability of EVs to access the central nervous system and support the concept that clinically manufactured MSC-derived EVs may serve as a practical platform for delivery of therapeutic nucleic acids to the brain.

The observed reduction in amyloid plaque burden is consistent with extensive genetic evidence supporting Bace1 as a key regulator of amyloidogenesis. Bace1 catalyzes the rate-limiting step in amyloidogenic APP processing, and genetic deletion of Bace1 in APP transgenic models effectively abolishes Aβ generation and plaque formation. Our findings suggest that EV-mediated siRNA delivery could achieve a degree of Bace1 suppression sufficient to alter disease-associated pathology without requiring complete target ablation; albeit future studies are needed to evaluate functional impact of EV-based therapeutics in this model.

Interestingly, the effects observed in our study appeared more pronounced in larger, high-intensity plaques than in smaller lesions. Although the current study was not designed to investigate mechanisms underlying plaque dynamics, this pattern may provide insight into the biological role of Bace1 during plaque maturation and growth. Previous studies have demonstrated that dystrophic neurites surrounding amyloid plaques exhibit elevated Bace1 expression and that local Bace1 accumulation contributes to ongoing amyloidogenic processing in the plaque microenvironment^16,28^. Suppression of Bace1 may therefore preferentially limit plaque expansion by reducing continued Aβ production at sites of established pathology. Under this model, smaller plaques that have already formed may be less sensitive to intervention over the treatment interval examined, whereas larger plaques dependent on sustained local Aβ generation may exhibit greater responsiveness. Future studies incorporating longitudinal analyses of plaque development and biochemical measurements of soluble and insoluble Aβ species will be necessary to further evaluate this possibility.

The translational relevance of the EV platform is strengthened by recent clinical experience with engineered EV therapeutics. We recently reported the first-in-human evaluation of MSC-derived engineered EVs/exosomes carrying Kras^G12D^-specific siRNA (iExosomes) in patients with metastatic pancreatic cancer^26^. Specifically, we evaluated biodistribution of IV administered labeled EVs in non-human primates and reported on brain localization of exogenously administered EVs. This study^26^ demonstrated a favorable safety profile, repeated systemic administration, target engagement, and evidence of biological activity in patients. Although the disease context differs substantially from Alzheimer’s disease, the clinical experience provides important validation of the underlying EV manufacturing process, product characterization, and systemic delivery approach. The current findings therefore extend the potential applicability of clinically translatable engineered EV platforms beyond oncology and into neurodegenerative disease.

Several limitations of the present study should be acknowledged. First, this work was designed primarily as a feasibility study and was not powered to evaluate functional outcomes. While reductions in plaque burden provide evidence of biological activity, future studies should incorporate behavioral assessments, learning and memory testing, and comprehensive neuroinflammatory profiling to determine whether the observed pathological changes translate into functional benefit. Similarly, quantitative analyses of soluble Aβ40 and Aβ42 species, APP processing intermediates, and downstream signaling pathways would provide additional evidence of target engagement and mechanism of action.

Second, these experiments were performed before broader recognition of the importance of parental transmission of the transgenes in the 5xFAD model. Subsequent studies have demonstrated that maternal versus paternal inheritance can significantly influence transgene expression, amyloid pathology, and disease progression in 5xFAD mice^29^. Because parental origin information was not available for the animals used in the current study, it remains possible that inheritance-related variability contributed to some of the observed heterogeneity. Future studies should incorporate controlled breeding strategies and documentation of transgene inheritance patterns to minimize this potential confounder.

Additional limitations include the relatively small cohort size and the absence of longitudinal imaging or molecular analyses that would permit assessment of treatment kinetics. Furthermore, although reductions in Bace1 protein and amyloid pathology were observed, the precise degree of target suppression achieved within specific neural cell populations remains unclear. Defining the relationship between EV dose, BACE1 suppression, and therapeutic effect will be important for future translational development.

In conclusion, this study establishes proof-of-concept feasibility for the use of engineered MSC-derived EVs as systemic carriers of Bace1-targeting siRNA in Alzheimer’s disease. The ability to deliver RNA cargo to the brain, achieve measurable target engagement, and reduce amyloid plaque burden following repeated systemic administration supports further development of EV-based therapeutic strategies for neurodegenerative disorders. Future studies incorporating larger cohorts, mechanistic analyses, and functional outcome measures will be necessary to determine the full therapeutic potential of this approach and to define optimal levels of Bace1 suppression for disease modification.

## Acknowledgments

We are grateful to Dr. Kathleen McAndrews and Michelle Kirtley for their guidance and assistance with data collection. Schemas in Figures 1B, 1H, and 2A were created with Biorender.com. The authors wish to also acknowledge the use of ChatGPT (https://chat.openai.com/) to assist with the writing of the manuscript.

## Funding

This work was supported by Black Rhino/Bosarge Family Trust Office to the Kalluri Laboratory at the University of Texas MD Anderson Cancer Center. Lyda Hill philanthropies inspired our efforts to use EVs as a delivery system for non-cancer related diseases.

## Conflict of Interests

RK and VSK are founders and equity owner of PranaX, Inc. All other authors declare no conflict of interest. RK and VSK are Members of the Editorial Board of Extracellular Vesicle.

## Supplementary Information

### Supplementary Methods

#### Flow cytometry analyses of EVs

EVs were characterized by flow cytometry for EV and MSC markers as previously described (see also Supplementary Text). Briefly, 2×10^10^ EVs were resuspended in 200 μL of PBS and treated with 15 μL of 3μm aldehyde-sulfate beads (Invitrogen; Cat. No. A2347389). The EV–bead mixture was incubated at room temperature for 15 min with gentle mixing on a tube rotator. Following incubation, 600 µL of 1x PBS was added, and the suspension was mixed overnight at 4°C. The following day, 400 µL of 1M glycine was added, and the mixture was incubated for 1 hour at room temperature with gentle mixing followed by centrifugation at 8,000*g* for 1 min, and subsequent pellet resuspension in 10% BSA and incubated for 45 min at room temperature. Subsequently, EV–bead complexes were incubated for 30 min at room temperature on a tube rotator with primary antibodies diluted in 2% BSA. The following antibodies were used at a 1:10 dilution: anti-CD47 (eBioscience; Cat. No. 14-0479), anti-CD63 (BD Biosciences; Cat. No. 556019), anti-CD81 (BD Biosciences; Cat. No. 555675), anti-CD29 (BioLegend; Cat. No. 303001), anti-CD90 (BioLegend; Cat. No. 328101), and mouse IgG1 isotype control (BD Biosciences; Cat. No. 555746). Anti-CD9 (MilliporeSigma; Cat. No. SAB4700092) was used at a 1:20 dilution. Following primary antibody incubation samples were washed with 200 µL of 2% BSA and centrifuged at 8,000*g* for 1 min. The pellet was then incubated with secondary antibody (anti-mouse Alexa fluor 488, Invitrogen; Cat. No. A21202) diluted in 2% BSA for 1 hour at room temperature on a tube rotator. Finally, EV–bead complexes were washed four times with 200 µL of 2% BSA, resuspended in 300 µL of 2% BSA, and analyzed using an LSR Fortessa flow cytometer.

#### In vitro BACE1 Targeting: primer information

Murine *Bace1* qPCR primers used are: Forward – 5’-ACCATCCTTCCTCAGCAATAC-3’and Reverse – 5’-ATGATGACGGCTCCCATAAC-3’. Murine *Actn* qPCR primers used are: Forward – 5’-GGCTGTATTCCCCTCCATCG-3’, Reverse. – 5’- CCAGTTGGTAACAATGCCATGT-3’. Human BACE1 qPCR primers used are: Forward – 5’- CCATCCTTCCGCAGCAATA-3’, Reverse – 5’- CGTAGAAGCCCTCCATGATAAC-3’. Human *ACTN* qPCR primers used are: Forward – 5’- CATGTACGTTGCTATCCAGGC -3’, Reverse. – 5’- CTCCTTAATGTCACGCACGAT -3’.

#### Biodistribution studies

For *in vivo* bio-distribution studies, EVs were labeled with DiI dye as previously reported^23^ and were resuspended in 100 µL of PBS prior to IV administration via the retro-orbital plexus. DiI control was prepared similarly but omitting EVs. Mice were euthanized 6 hours and 12 hours post injection, and tissues were removed and fixed overnight in 4% paraformaldehyde before incubating in 30% sucrose for 48 hours. Sections (5 µm) were cut from tissues embedded in optimal cutting temperature (OCT) medium. Sections were stained with 0.25µg/mL DAPI in TBS for 5 minutes then mounted with coverslips using Fluoroshield (Sigma). Slides were imaged using Zeiss LSM800confocal microscope.

## Supplementary Figure Legends

**Supplementary Figure 1.**
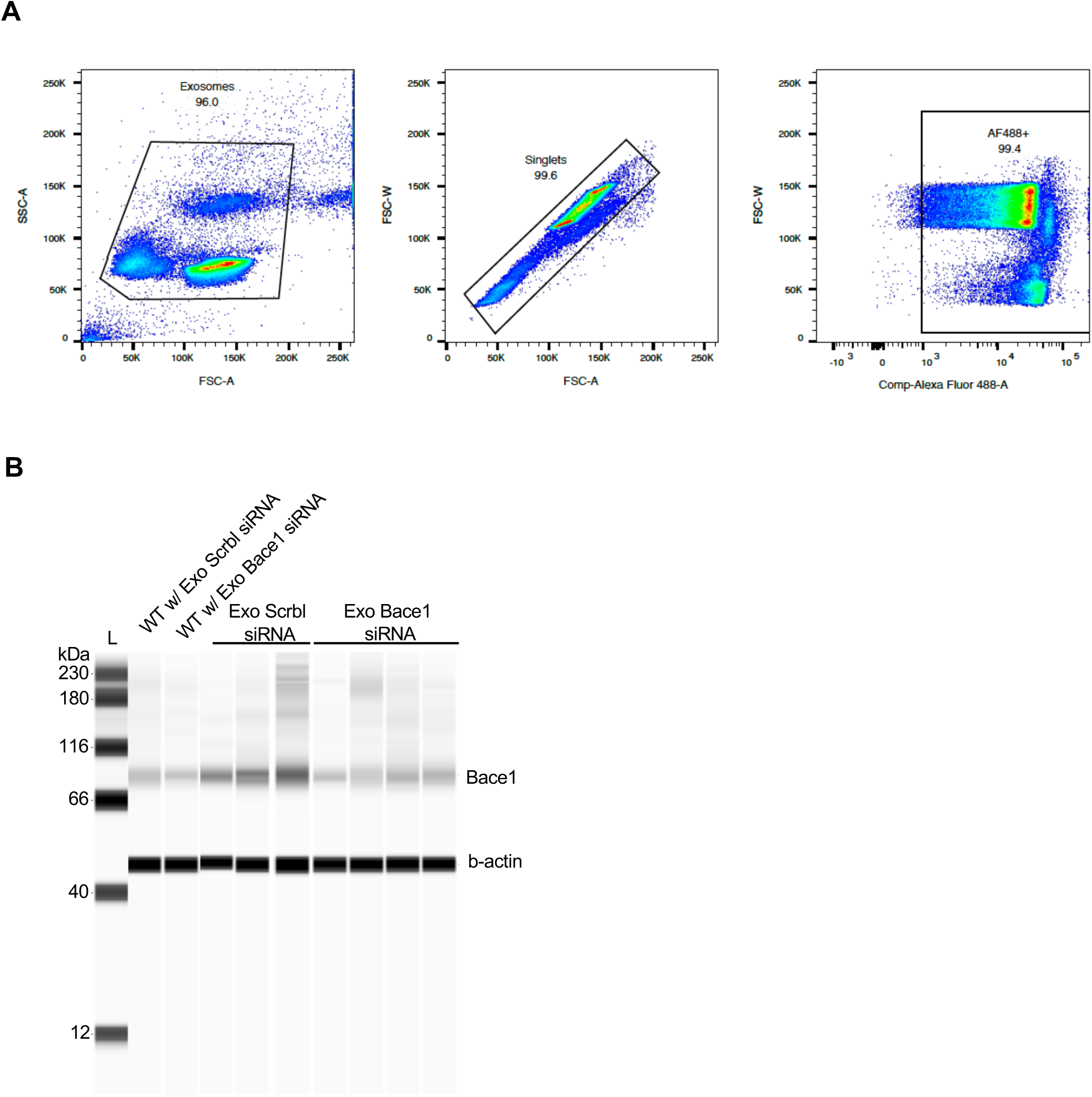
Supporting data for Figure 1D and 3F. **A.** Flow cytometry gating strategy employed for data presented in Figure 1D. **B**. Full length run of electrophoretic migration of brain lysate proteins assayed for Bace1 and β-actin **shown in** Figure 3. L: ladder. kDa: kilodalton.

**Supplementary Figure 2.**
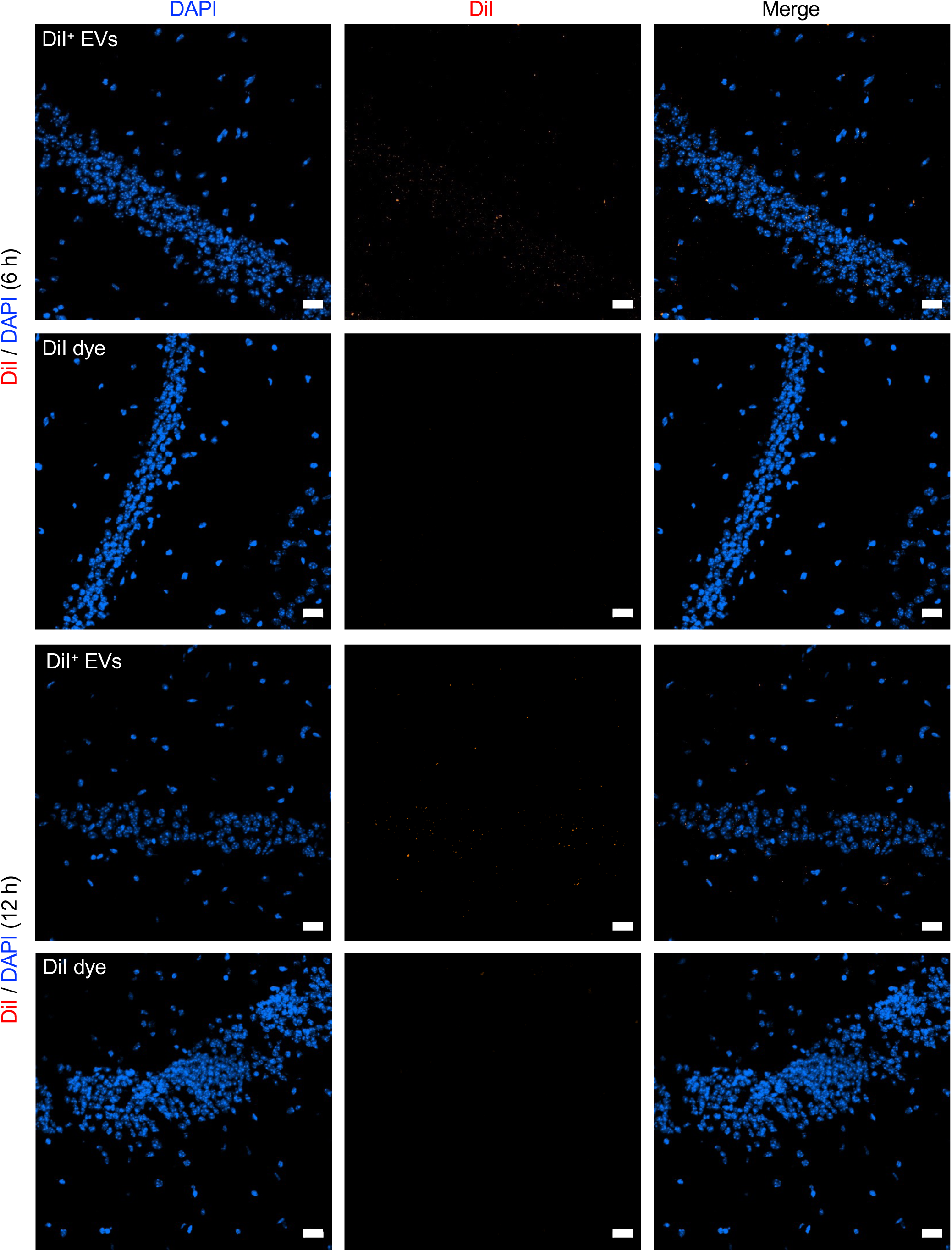
Supporting data for Figure 1H. Individual channels shown for brain sections evaluated for DiI signal 6 hours or 12 hours after either DiI^+^ EVs or DiI dye only IV administration to mice. Data also **shown in** Figure 1H. Scale bar: 20 μm. DAPI: nuclear stain.

**Supplementary Figure 3.**
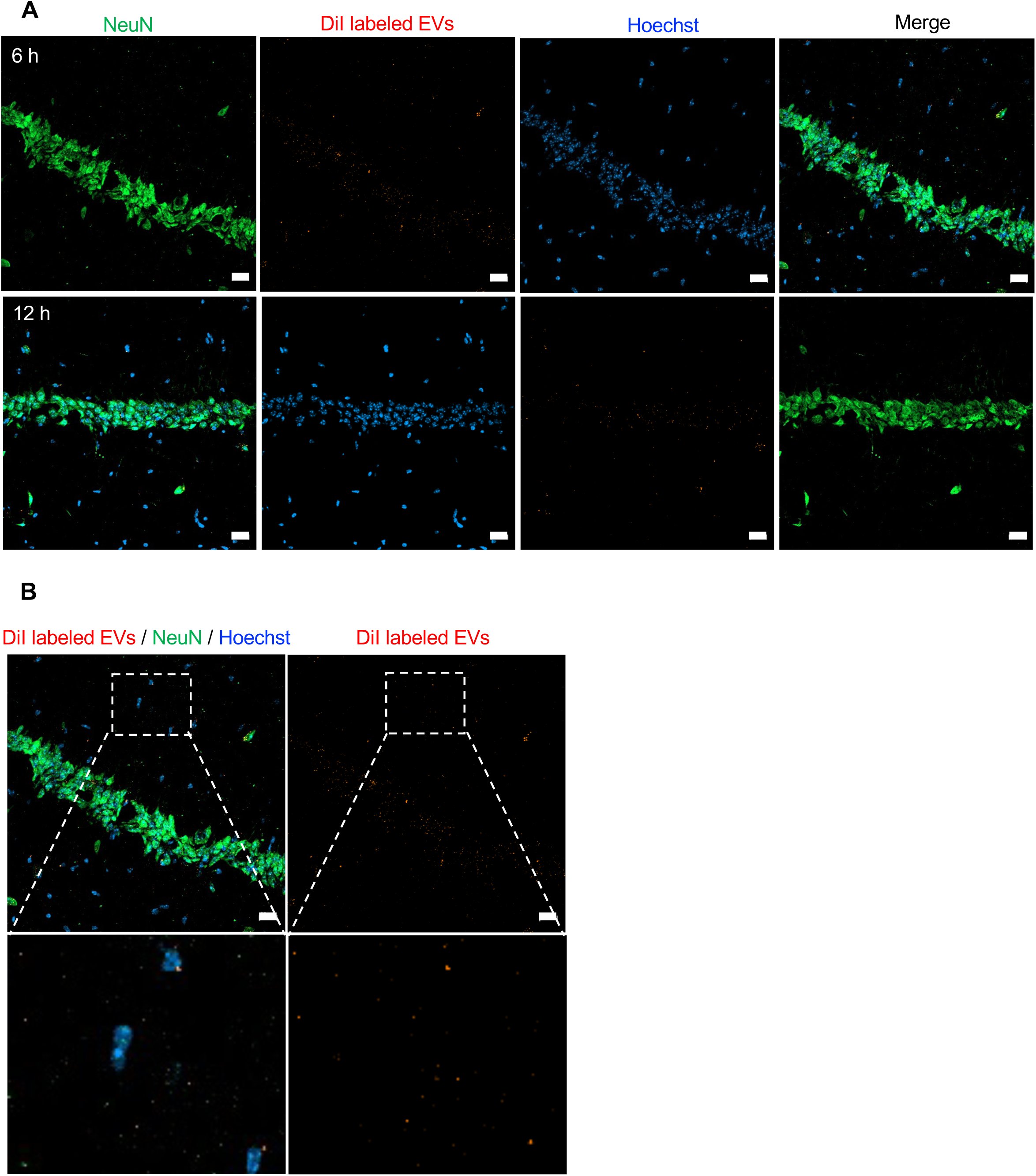
Supporting data for Figure 1I. **A**. Individual channels shown for brain section for DiI, NeuN, and nuclear stain (Hoechst) at 6 hours and 12 hours post EV administration. **Data also shown in** Figure 1I **for 6 hours**. Scale bar: 20 μm. **B.** Representative image of brain section for DiI, NeuN, and nuclear stain (Hoechst) with digital zoom at 6 hours post EV administration in Neu-negative field of view (left). Single channel for DiI labeled EVs also shown (Right). Scale bar: 20 μm.

